# Human pulmonary neuroendocrine cells respond to House dust mite extract with PAR-1 dependent release of CGRP

**DOI:** 10.1101/2024.09.09.612124

**Authors:** Ritu Mann-Nüttel, Shivani Mandal, Marie Armbruster, Lakshmi Puttagunta, Paul Forsythe

## Abstract

**Background:** Pulmonary neuroendocrine cells (PNEC) are rare airway epithelial cells that have recently gained attention as potential amplifiers of allergic asthma. However, studying PNEC function in humans has been challenging due to a lack of cell isolation methods and little is known about human PNEC function in response to asthma relevant stimuli. Here we developed and characterized an *in vitro* human PNEC model and investigated the neuroendocrine response to extracts of the common aeroallergen house dust-mite. (HDM).

**Methods:** PNEC enriched cultures were generated from human induced pluripotent stem cells (iPNEC) and primary bronchial epithelial cells (ePNEC). Characterized PNEC cultures were exposed to HDM extract, a volatile chemical odorant (Bergamot oil), or the bacterial membrane component, lipopolysaccharide (LPS) and neuroendocrine gene expression and neuropeptide release determined.

**Results:** Both iPNEC and ePNEC models demonstrated similar baseline neuroendocrine characteristics and a stimuli specific modulation of gene expression. Most notably, exposure to HDM but not Bergamot oil or LPS, lead to dose dependent induction of the CGRP encoding gene, CALCB, and corresponding release of the neuropeptide. HDM induced CALCB expression and CGRP release could be inhibited by a protease activated receptor 1 (PAR1) antagonist or protease inhibitors and was mimicked by a PAR1 agonist.

**Conclusions:** We have characterized a novel model of PNEC enriched human airway epithelium and utilized this model to demonstrate a previously unrecognized role for human PNEC in mediating a direct neuroendocrine response to aeroallergen exposure and highlighting CGRP production by these cells as a potential therapeutic target in allergic asthma.

## Introduction

Pulmonary neuroendocrine cells (PNEC) are rare neuroepithelial cells found in clusters at airway branch points and as solitary cells scattered throughout the airways [1]. They contain dense core-vesicles packed with various neuropeptides including Calcitonin gene-related peptide (CGRP), Gamma-aminobutyric acid (GABA) and serotonin [2-5]. Innervated PNEC initiate sensory signal transduction from the airway epithelium to the central nervous system (CNS) via the vagus nerve [6]. Studies suggest PNEC act as chemosensory sentinels of the inhaled environment responding to diverse stimuli including volatile chemicals [7]. Recently, PNEC gained attention as potential amplifiers of the allergic airway response [8]. Studies in mouse models of asthma suggest that PNEC derived CGRP can stimulate Innate lymphoid cells 2 (ILC2) [8] cells to elicit downstream type 2 immune responses, enhancing airway inflammation [8]. In humans, asthma is associated with PNEC hyperplasia [8] and it now seems possible that the mediator content and responsiveness of PNEC contributes to the development and severity of the disease. Thus, PNEC may represent novel target cells for asthma therapeutics. However, while data from murine models are encouraging, little is known regarding human PNEC function and response to asthma relevant stimuli, including allergens. This knowledge gap exists largely because the cells make up such a small proportion (0.04%) [9] of the lung epithelium and there are currently no suitable methods for isolation. To address this issue, we utilized two *in vitro* human PNEC models. The first (iPNEC) was derived from induced pluripotent stem cells (iPSC) as described by Hor et al [10]. The second was a novel model of PNEC enriched human airway epithelium (ePNEC) derived from primary bronchial/tracheal epithelial cells (HBEC) that has not previously been characterized. We aimed to determine ePNEC and iPNEC expression of canonical neuroendocrine markers and assess cultured human PNEC responses to representative stimuli from the inhaled environment: House dust mite (HDM) as an allergen, LPS as bacterial stimulus, and Bergamot oil as a volatile chemical often contained in perfumes. Using this approach we demonstrate, for the first time, that human PNEC respond to challenge by extracts of HDM to produce the immunomodulatory peptide CGRP. We also provided mechanistic insight identifying that the HDM response is dependent on protease activated receptor 1 (PAR1). This study suggests a previously unrecognized role for human PNEC in mediating a direct neuroendocrine response to allergen exposure.

## Results

### Confirmation of iPNEC and ePNEC development

We differentiated iPNECs from iPSCs derived from adult skin fibroblasts expressing typical iPSC markers Homeobox Transcription Factor NANOG and POU Class 5 Homeobox 1 (POUF5F1). ePNECs were differentiated from HBEC expressing Keratin 5 (KRT5) and Tumor protein 63 (TP63) (**Fig. S1A**). iPNEC and ePNEC enrichment was induced at ALI in the presence of Notch signaling inhibitor 3tert-Butyl(2S)-2-((2-(3,5-diflurophenyl) acetyl) amino) propanoyl)amino)-2-phenylacetate (DAPT) at 10 μM (**Fig. S1B, E**). At day 30 of culture we found expression of the pro-neural transcription factor ASCL1 (Achaete-Scute Family BHLH Transcription Factor 1), a known PNEC cell fate factor [11] and Enolase 2 (**Fig.S1 C**), and at day 60 of culture, we found expression additional neuroendocrine markers Chromogranin A (CHGA), Roundabout receptor 2 (ROBO2) and Synaptophysin (SYP) (**Fig. 1A**). These markers indicate that PNECs developed in both cultures. In quantifying the proportion of PNEC in the cultures, expression of CHGA and SYP was determined as a percentage of the mean fluorescent intensity (MFI) of the respective marker normalized to the expression of a nuclear marker (Hoechst dye) at day 60 of culture. This identified 23.4 ± 3.7 and 25.9 ± 2.3 % CHGA^+^ cells in the iPNEC and ePNEC culture, respectively (**Fig. 1B, left**). The proportion of SYP^+^ cells in iPNEC and ePNEC cultures at day 60 was 19.9 ± 2.1 and 26.6 ± 1.6 % respectively (**Fig. 1B and C**). Further, we performed single cell RNA sequencing of ePNEC culture and identified two PNEC clusters by expression of ASCL1 or CHGA, among other markers (**Fig.S1 F**) representing 21% of the cells sequenced. In addition, flow cytometry experiments for CHGA staining confirmed presence of CHGA^+^ cells in our ePNEC cultures (**Fig.S1 D**). Other cell types identified in the ePNEC culture system were goblet cells, club cells, basal cells, ciliated epithelial cells and myofibroblasts (**Fig. 1D, S1G**).

**Fig 1.**
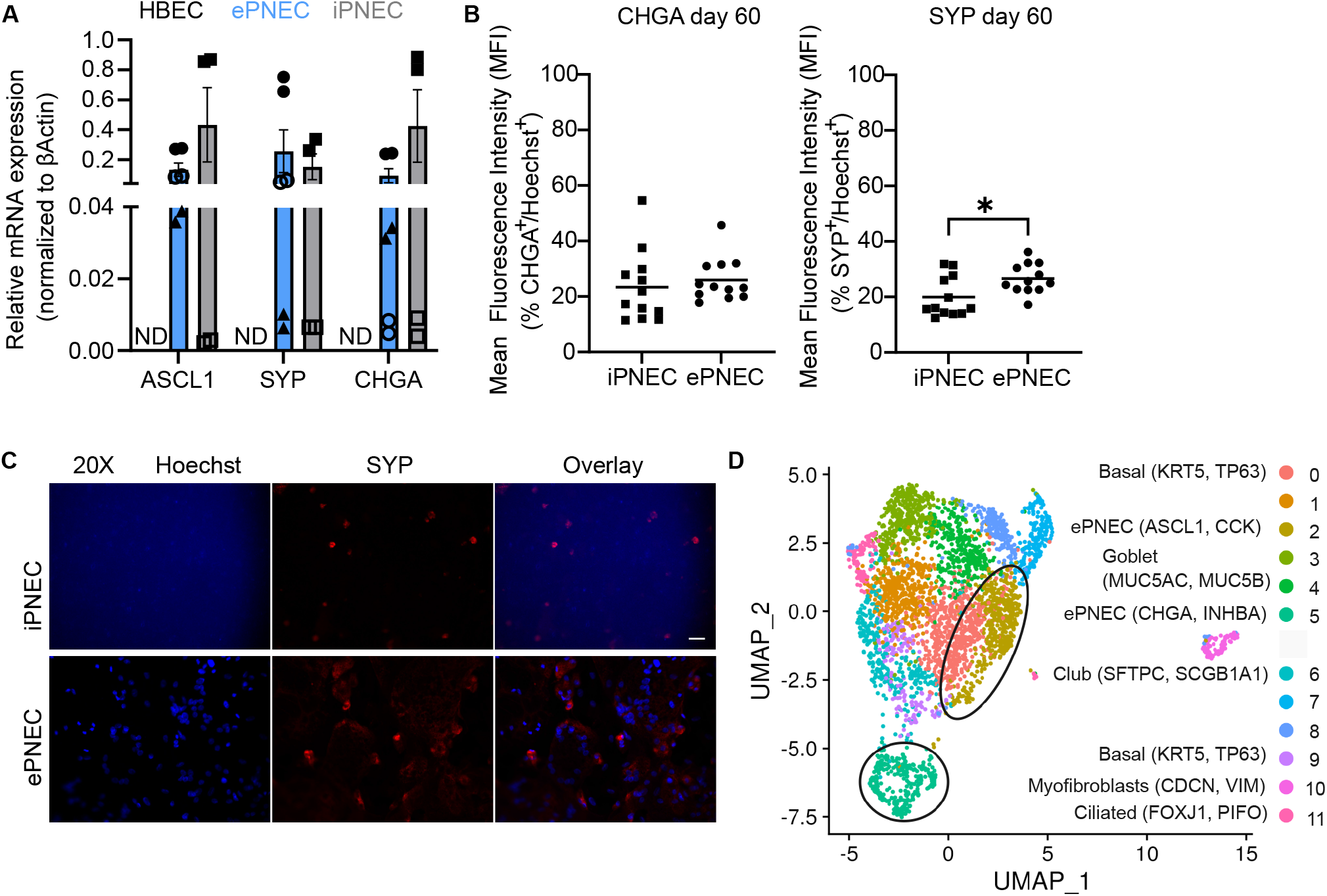
Characterization of iPNEC and ePNEC cultures. **A** Quantitative RT-PCR showing expression of characteristic PNEC markers in human cultured iPSC and -HBEC-derived iPNEC (□ female, 22; ◼ male, 32) and ePNEC (● male, 52; ▲ male, 56; ○ female, 55) and control HBEC at day 60 in ALI culture, respectively. **B** Mean fluorescent intensity (MFI) of CHGA^+^/Hoechst^+^ cells (left) and SYP^+^/Hoechst^+^ cells [34] at day 60. Data is representative of four different culture plate wells per group for one biological donor for each cell type (ePNEC male, 52; iPNEC male, 32). **C** Representative IF images show SYP^+^ cells and nuclei are counterstained with Hoechst. 20X magnification and scale bars at 100 μm. **D** Single cell RNA sequencing identifies two ePNEC clusters in 60-day old ePNEC differentiated cells. Each dot represents one well and data shown for mean ± SEM. Mann-Whitney test performed for A, B; *<0.05,**<0.01, ***<0.001, ****<0.0001. Abbreviations: ASCL1 -Achaete-Scute Family BHLH Transcription Factor 1; CHGA – Chromogranin A ; ePNEC – epithelial-derived pulmonary neuroendocrine cells; iPNEC – iPSC-derived pulmonary neuroendocrine cells; iPSC – induced pluripotent stem cells; SYP – Synaptophysin.

### iPNEC and ePNEC cultures have neuroendocrine phenotype and function

Recently, Kuo et al. assessed the potential expression of all peptidergic genes annotated in the mouse genome in *ex vivo* sorted murine PNECs [12]. We selected 16 genes from their study to evaluate expression in human cultured PNEC (**Fig. 2A**). iPNEC and ePNEC cultures expressed an identical profile of neuroendocrine markers with 12 of the 16 genes assessed expressed constitutively in both cultures, none of which were detectable in iPSC or HBEC. The GABA producing enzyme, GAD67, Gastrin releasing peptide GRP or genes responsible for CGRP production (CALCA and CALCB) were not expressed in either culture system at baseline. The expressed genes included is the PNEC driving transcription factor ASCL1, the receptor ROBO2, neuropeptides Proenkephalin (PENK), Somatostatin (SST), Pituitary adenylcyclase-activated peptide (ADCYP1), Corticotropin releasing hormone (CRH), and neurotransmitters synaptic vesicle protein 2 (SV2), Cocaine and amphetamine-related transcript (CARTPT), Choline O-Acetyltransferase (CHAT) encoding an enzyme that makes Acetylcholine, Dopa Decarboxylase (DDC), Palmitoyl-Protein Thioesterase 1 (PPT1), Synaptotagmin 1 (SYT1), a master regulator for neurotransmitter release, and Enolase 2 (ENO2), a biomarker for neuroendocrine-related diseases, and the granin gene CHGA. These results indicate that our developed ePNEC model has an overall similar pattern of neuroendocrine related gene expression to the published iPNEC model [10], however certain genes such as SST are expressed at different relative levels between iPNEC and ePNEC. Notably, control HBEC cultured at ALI for 60 days without DAPT and other PNEC promoting factors did not express any typical PNEC markers (**Fig. 2B**). Furthermore, we observed inter-individual differences between our 3 donors regarding the levels of expressed neuroendocrine markers, suggesting that intrinsic factors may influence the expression level and/or number of PNEC in cultures depending on the donor (**Fig. 2B**).

**Fig 2.**
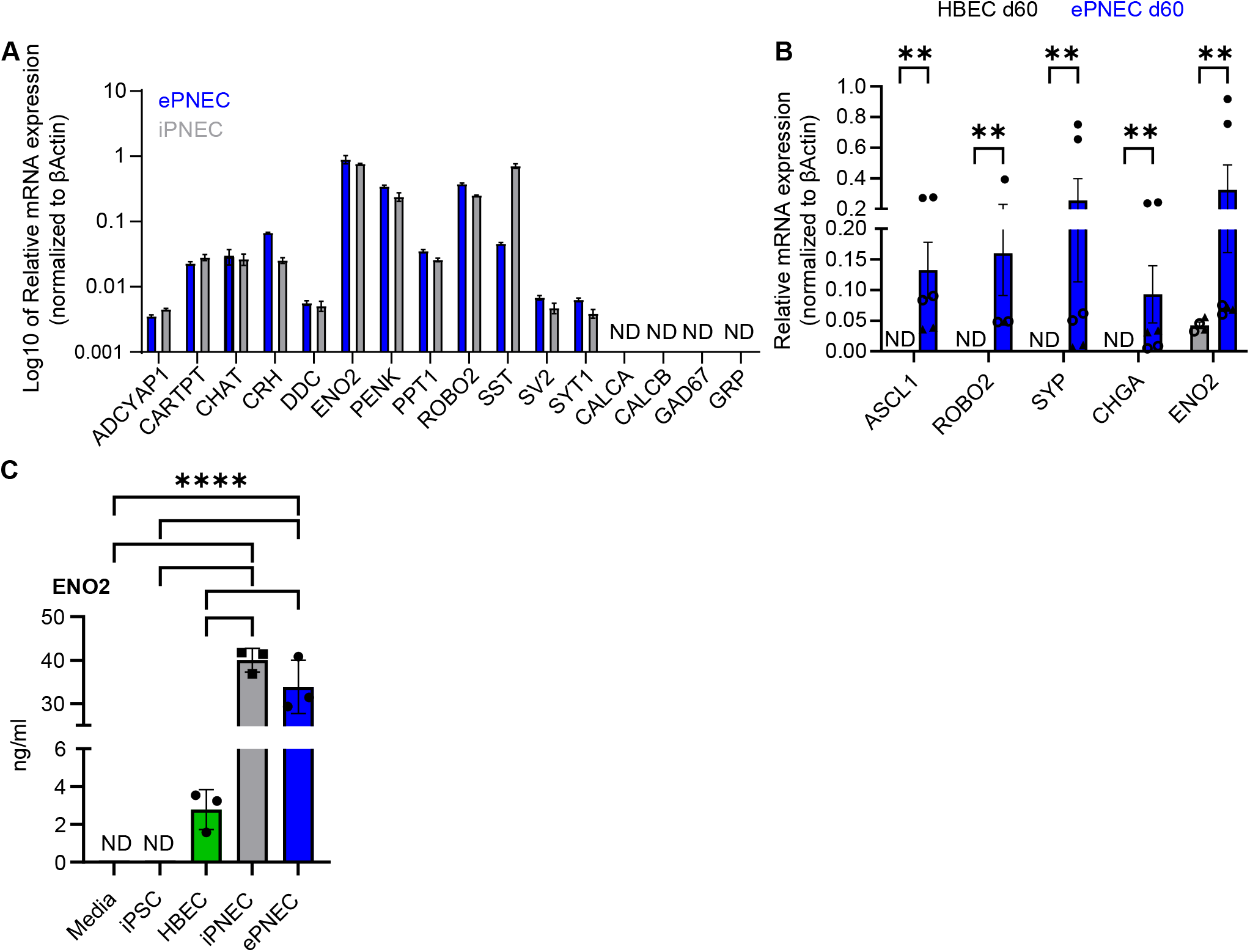
Expression of neuroendocrine markers in iPNEC and ePNEC cultures. **A, B** Quantitative RT-PCR showing expression of neuroendocrine markers in **A** human cultured iPSC and -HBEC-derived iPNEC (male, 32) and ePNEC (male, 52). **B** control HBEC and ePNEC (• male, 52; ▴ male, 56; ○ female, 55) at day 60 in ALI culture, respectively. **C** Enzyme-linked immunosorbent assay (ELISA) data showing secreted levels of ENO2 for one donor per cell type (iPNEC, male, 32; HBEC and ePNEC male, 52) in the basolateral culture media. Each dot represents one well and data shown for mean ± SEM. Mann-Whitney test for A, B, One-way ANOVA for C, D; *<0.05, **<0.01, ***<0.001, ****<0.0001. Abbreviations: ADCYAP1-Adenylate cyclase activating polypeptide 1; CALCA – Calcitonin-related polypeptide alpha; CALCB – Calcitonin-related polypeptide beta; CGRP – Calcitonin-gene related peptide; CARTPT – Cocaine and amphetamine regulated transcript; CHAT – Coline O-Acetyltransferase; CRH – Corticotropin releasing hormone; DDC -Dopa Decarboxylase; ENO2 – Enolase 2; ePNEC – epithelial-derived pulmonary neuroendocrine cells; GAD67 -Glutamic acid decarboxylase 67; GRP – Gastrin releasing peptide; HBEC – human bronchial epithelial cells; iPNEC – iPSC-derived pulmonary neuroendocrine cells; PENK – Proenkephalin; PPT1 – Palmitoyl-protein thioesterase 1; ROBO2 – Roundabout Guidance receptor 2; SST – Somatostatin; SV2 – Synaptic vesicle protein 2; SYP – Synaptophysin; SYT1 – Synaptotagmin 1.

We also confirmed Enolase secretion, first described by Hor et al. [13] as a characteristic of PNEC cultures, with increased enolase in the basolateral transwells of both iPNEC (40.0 ± 1.6) and ePNEC (33.9 ± 3.5 ng/ml) compared to HBEC (2.8 ± 0.6 ng/ml) (**Fig. 2C**).

### Gene expression in cultured human PNEC is altered in response to stimuli

We assessed changes in expression of 12 genes related to neuroendocrine activity in response to LPS, HDM and the volatile chemical Bergamot oil (**Fig. 3A**). Of the genes assessed 10 were expressed constitutively in naïve iPNEC and ePNEC cultures. LPS induced significant expression changes for all 10 genes in iPNEC (up-regulated ENO2, ADYAP1, CARTPT, CHAT, PENK, ROBO2, SST; down-regulated CRH, PPT1, SYT1), and for 6 genes in ePNECs (up-regulated ENO2, CHAT, PENK, PPT1, ROBO2, SST). Bergamot oil, a volatile chemical known to release serotonin and CGRP from primary PNECs [7], had a significant effect on the expression of 8 genes in iPNECs (up-regulated ENO2, CARTPT, CHAT, PENK; down-regulated ADCYAP1, CRH, SST, SYT1) and in 7 genes in ePNEC (up-regulated ENO2; down-regulated ADCYAP1, CARTPT, CHAT, PENK, ROBO2, SYT1).

**Fig 3.**
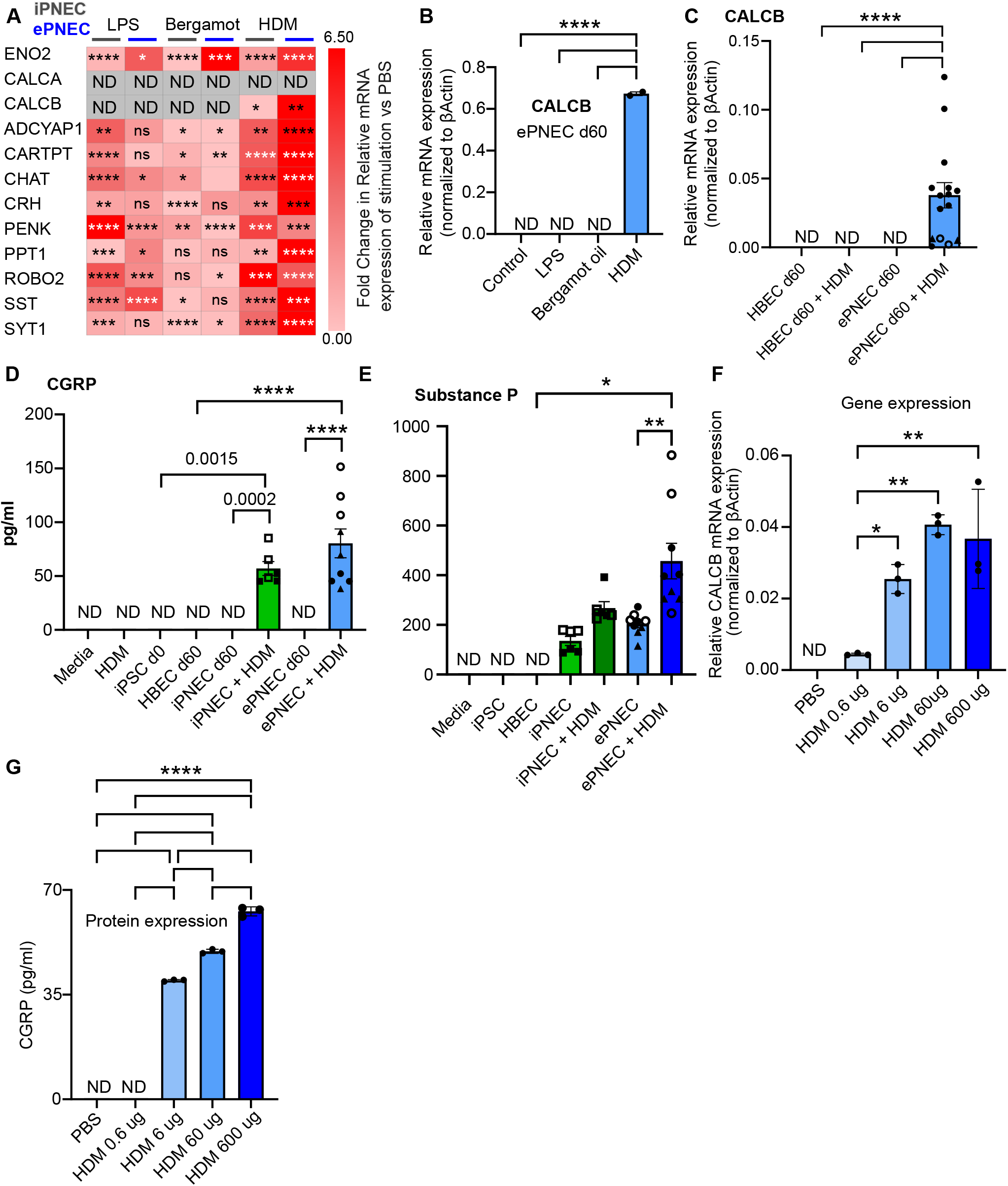
HDM, but not LPS or Bergamot, induce CGRP expression in iPNEC and ePNEC. **A** Heatmap showing RT-PCR results for fold change of expression of stimulated iPNEC (male, 32) and ePNEC (male, 52) with LPS, Bergamot oil and HDM for 2h relative to control stimulated cells with PBS only and normalized to internal control β-Actin. Grey color signifies no expression **B** qRT-PCR of CALCB in control (PBS), 50 μL HDM (1200 μg/ml), LPS (1ng/μl) and Bergamot oil (1:1000, PBS) stimulation of ePNEC (male, 52) for 2h. **C** qRT-PCR of CALCB in control (PBS) and 2h 50 μL HDM (1200 μg/ml) stimulated ePNEC and control HBEC cultured cells (● male, 52; ▲ male, 56; ○ female, 55) at ALI for 60 days. **D** CGRP ELISA of basolateral supernatant of iPNEC (□ female, 22; ◼ male, 32) and ePNEC (● male, 52; ▲ male, 56; ○ female, 55) stimulated with 50 μL HDM (1200 μg/ml in PBS) for 6h. **E** Substance P for naïve and 6h HDM stimulated iPNEC (□ female, 22; ◼ male, 32) and ePNEC (● male, 52; ▲ male, 56; ○ female, 55) in the basolateral culture media. **F, G** qRT-PCR of the CALCB gene (**F**) and CGRP ELISA for basolateral supernatant (**G**) after stimulation of ePNEC (male, 52) at different HDM dilutions (50 μl of 12, 120, 1200 and 12000 μg/mL corresponding to 0, 0.6, 6, 60, and 600 μg total protein in PBS) and PBS control. Each dot represents one well and data shown for mean ± SEM. Multiple unpaired t-tests for **A**, Kruskal-Wallis test for **C**, One-way ANOVA for **B, F, G**, unpaired t-test for **D;** *<0.05, **<0.01, ***<0.001, ****<0.0001. Abbreviations: ADCYAP1 – Adenylate cyclase activating polypeptide 1; ASCL1 -Achaete-Scute Family BHLH Transcription Factor 1; CALCA – Calcitonin-related polypeptid2e alpha; CALCB – Calcitonin-related polypeptide beta; CGRP – Calcitonin-gene related peptide; CARTPT – Cocaine and amphetamine regulated transcript; CHAT – Coline O-Acetyltransferase; CHGA – Chromogranin A ; CRH – Corticotropin releasing hormone; ENO2 – Enolase 2; ePNEC – epithelial-derived pulmonary neuroendocrine cells; HDM – House dust mite; iPNEC – iPSC-derived pulmonary neuroendocrine cells; iPSC – induced pluripotent stem cells; LPS – Lipopolysaccharide; PENK – Proenkephalin; PPT1 – Palmitoyl-protein thioesterase 1; ROBO2 – Roundabout Guidance receptor 2; SST – Somatostatin; SYP – Synaptophysin; SYT1 – Synaptotagmin 1.

### House dust mite induces CALCB expression and CGRP release from PNEC

Exposure to HDM extract (*Dermatophagoides pteronissinus*, 1200 μg/mL) significantly increased the expression of 11 genes in both iPNEC and ePNEC. tStrikingly, HDM, but not LPS or Bergamot oil, induced expression of the CGRP gene, CALCB, in both iPNEC and ePNEC (**Fig.3 A, B**). Expression of CALCB was not observed in unstimulated ePNEC, or control HBEC cultured at ALI for 60 days left naïve or exposed to HDM (**Fig. 3C**). PNEC markers ASCL1, SYP and CHGA maintained stable expression (**Fig.S2 A, B**). ELISA confirmed release of CGRP in the supernatant of HDM challenged iPNEC (47.5 ± 2.4 pg/mL) and ePNEC (30.2 ± 0.6 pg/mL) but was not detectable in unstimulated cells or HDM exposed HBEC (**Fig. S2 C**). To further characterize the response to HDM we stimulated ePNEC with a range of allergen dilutions (50 μl of 12, 120, 1200 and 12000 μg/mL corresponding to 0, 0.6, 6, 60, and 600 μg total protein in PBS) and identified a dose dependent response for CALCB mRNA induction at 2h (**Fig. 3F**) and CGRP protein expression at 6h. ranging from 0.6 μg (no CGRP), 6 μg (39.77 ± 0.05 pg/ml CGRP), 60 μg (49.50 ± 0.10 pg/ml CGRP) to 600 μg (61.85 ± 0.19 pg/ml CGRP) (**Fig. 3G**).

The neuroendocrine response to HDM was not restricted to CGRP. In Contrast to CALCB, the Substance P encoding gene, Preprotachykinin A (PPT1), was expressed at baseline in both PNEC cultures (**Fig. 2A**), with a corresponding release of the neuropeptide into the basolateral supernatant of naive iPNEC and ePNEC cultures (135.5 ± 18.5 pg/ml and 203.7 ± 14.8 pg/ respectively). HDM stimulation of either iPNEC or ePNEC led to a significant increase in PPT1 expression (**Fig. 3A**) and substance P release (268.2 ± 25.4 pg/ml (p=0.0290) and 457.3 ± 71.7 pg/ml (p=0.0039)), respectively. (**Fig. 3E**). We also compared the neuropeptide response to that of the typical epithelial derived cytokines [14], Interleukin-8 (IL-8) and Granulocyte macrophage-colony stimulating factor (GM-CSF) in ePNEC cultures. Both cytokines were present in culture supernatants at baseline, but unlike CGRP and Substance P cytokine levels were not increased 6 hours after HDM exposure. However, both IL-8 and GM-CSF were significantly increased 24 hours following HDM exposure (**Fig.S2 D**).

### CALCB induction by HDM is Protease activated receptor 1 (PAR1)-mediated

We next investigated the mechanism underlying HDM induced CGRP production. We assessed PNEC expression of receptors implicated in allergen sensing [15, 16] through a re-analysis of published single cell transcriptomics data of iPNECs (GSE146990, [13]) and our own ePNEC data. Notably, we found that the receptor PAR1 encoded by the gene F2R was expressed in a subpopulation of iPNECs and ePNEC, whereas expression of other PAR family members was either absent or expressed at much lower levels . In addition, PAR1 expression is markedly higher in iPNEC and ePNEC clusters compared to the other cell types in the cultures (**Fig. 4A**). We also performed immunohistochemical co-staining of SYP and PAR1 to confirm PAR1 expression on PNEC cells (**Fig. 4B**). To test the hypothesis that PAR1 is required for CALCB induction after HDM stimulation we used the PAR1 antagonist Vorapaxar and the PAR1 agonist TFLLR-NH2. TFLLR-NH2 mimicked the effect of HDM in inducing CALCB and CGRP expression while Vorapaxar blocked this response (**Fig. 4C, D**). In contrast, the PAR2 inhibitor FSLLRY did not significantly affect CALCB expression during HDM stimulation (**Fig.S3 B**). *In vivo*, CGRP half-life is limited due to the sensitivity of the peptide to protease degradation. Using PAR1 agonist stimulation we demonstrated that CGRP levels were not significantly altered by the presence of a broad-spectrum protease inhibitor in the basolateral chamber, suggesting that endogenous proteases were unlikely to influence detection of the neuropeptide in our culture system (**Fig.S2 E**).

**Fig 4.**
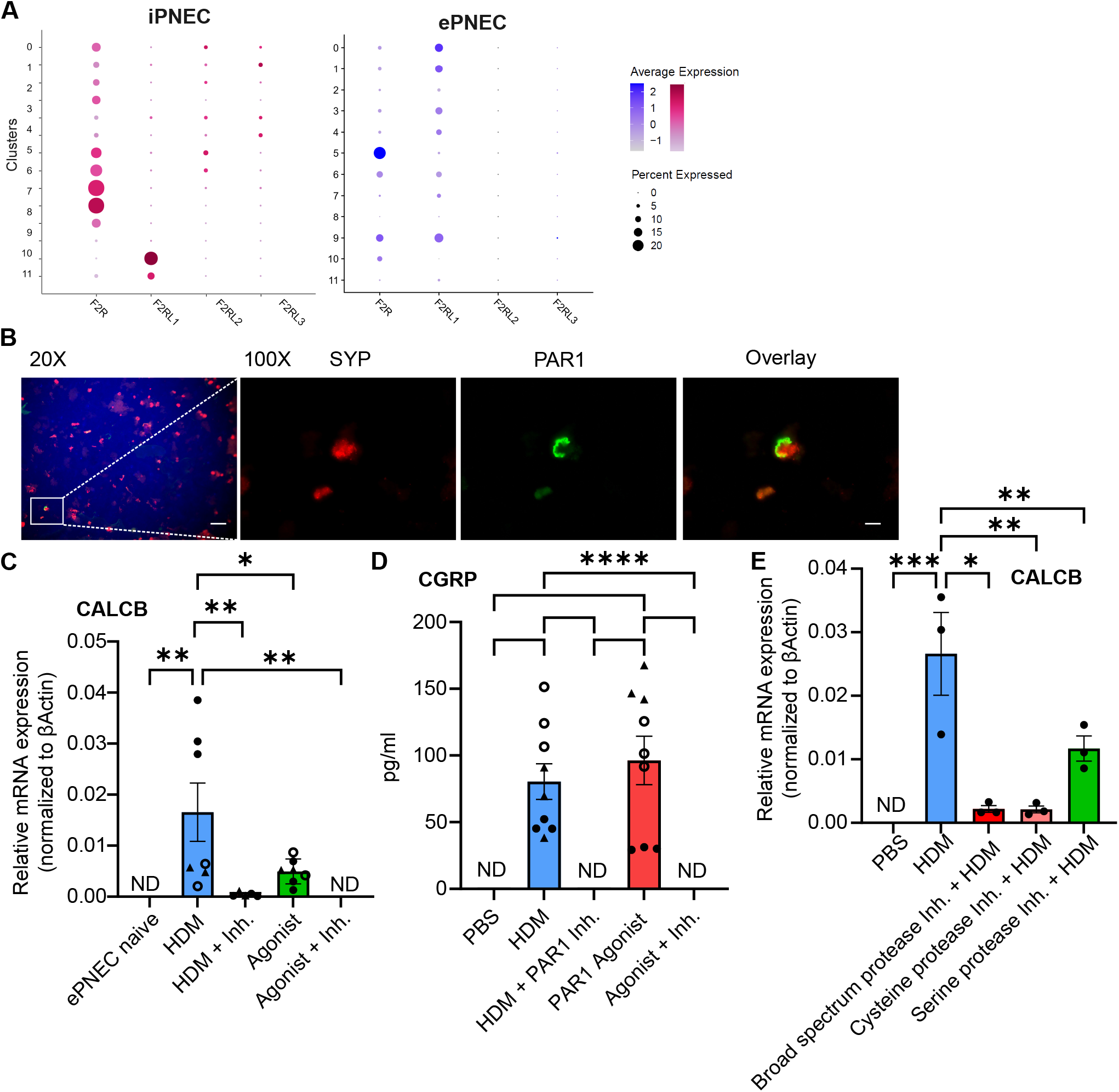
CGRP release after HDM stimulation is PAR1-dependent. **A** Single-cell transcriptomics data of iPNECs (GSE146990, Hor et al.) and ePNEC for PAR1 (F2R), PAR2 (F2RL1), PAR3 (F2RL2) and PAR4 (F2RL3) genes. **B** SYP and PAR1 immunohistochemical co-staining in naive iPNEC (male, 32). 20X and 100X magnification and scale bars at 50 μm and 10 μm, respectively. **C, D** CALCB mRNA (**C**) and CGRP protein (**D**) expression in different ePNEC (● male, 52; ▲ male, 56; ○ female, 55) conditions after 2h: 50 µL PBS, HDM (1200 μg/ml), PAR1 inhibitor Vorapaxar (80uM) and PAR1 agonist TFLLR-NH2 (2uM). **E** CALCB mRNA RT-PCR expression in different ePNEC conditions after 2h: 50 µL PBS, HDM (1200 μg/ml), HDM co-incubation with protease inhibitors Chymostatin (broad spectrum, 10 μg/ml), PMSF (serine specific, 0.25 mM) and E-64 (cysteine specific, 10 μM). Inhibitor/agonist and or HDM added at the same time. Each dot represents one well and data shown for mean ± SEM. One-way ANOVA for C, D, E; *<0.05, **<0.01, ***<0.001, ****<0.0001. Abbreviations: Agonist – PAR1 agonist vorapaxar; ePNEC – epithelial-derived pulmonary neuroendocrine cells; Inh. – PAR1 inhibitor; iPNEC – iPSC-derived pulmonary neuroendocrine cells; HDM – House dust mite; PAR1 – Protease activated receptor 1; SYP – Synaptophysin.

To gain further mechanistic insight into which HDM extract component might be responsible for CALCB induction we repeated our stimulation experiments in the presence of various protease inhibitors. Our results indicate that CALCB mRNA is significantly reduced in the presence of the broad spectrum protease inhibitor Chymostatin, as well as the serine and cysteine specific protease inhibitors PMSF and E-64, respectively (**Fig. 4E**). Of note, the reduction in CALCB mRNA was less with PMSF as compared to the other inhibitors, suggesting a cysteine protease such as Der P1 [17, 18] may be involved in CALCB mRNA induction in ePNEC. However, the role of Der P1 could not be verified in experiments with recombinant protein, despite protease activity being confirmed with and without Dithiothreitol activation (**Fig.S3 C, D**). Taken together, these results suggest that HDM induced CALCB expression and subsequent release of CGRP by PNEC is PAR1-dependent.

## Discussion

Here we established two differentiation strategies for growing human PNEC that overcome the limitations of isolation from lung tissue and allow us to study cell function *in vitro*. The first model, iPNEC, was previously reported and characterized by Hor et al. [10, 13] and, using a different source of iPSC, we have confirmed their observations that this approach leads to cells expressing canonical PNEC markers including CHGA, SYP, ASCL1, Synaptic vesicle protein 2 (SV2) and ROBO2, and high expression of both SST and ENO2. We also demonstrated, for the first time, constitutive expression and release of the neuropeptide Substance P by these cells. Notably, based on evidence that PNEC could potentially be generated from primary HBEC [13] we also developed a second approach to generate neuroendocrine cells. We demonstrated that treatment with a combination of growth factors together with inhibition of NOTCH signaling in HBEC cells at ALI resulted in the generation of cells with neuroendocrine characteristics that were comparable in both proportion and expression profile of 12 neuroendocrine related genes as to those obtained from iPSC. Single cell RNA sequencing data from Hor et al. showed that iPNEC cultures contain ∼30% iPNEC and induced basal cells, respiratory smooth muscle-like cells and fibroblast-like cells, among some unidentified cells [13]. Our own sequencing experiment, showed that ePNEC cultures contain ∼21% PNEC with additional cells consisting largely of proximal airway epithelial cells (club cells, ciliated cells, basal cells, goblet cells). The ePNEC model thus appears to have a greater variety of airway epithelial cells than the iPNEC model better mimicking the natural microenvironment of PNEC in vivo. Further, the ePNEC model has the advantages of being a simpler process than iPNEC generation and offers the potential to study PNEC derived from the airways of both healthy individuals and those with pulmonary disease.

As one of the very few innervated chemosensory cells in the airway epithelium, PNEC have been suggested as key sentinels for the inhaled environment [19]. Primary human PNEC have been shown to release Adenosine triphosphate in response to succinate [20] and mouse PNEC released CGRP and GABA after activation with eosinophil extracellular traps [21] and in a model of allergic asthma [8]. A mouse model of asthma showed that mice lacking PNEC had reduced airway inflammation following ovalbumin sensitization [8], suggesting that endogenous mediators must be able to activate the secretion of neuroendocrine mediators in PNEC. However, there have been no previous reports of human *in vitro* differentiated PNEC response to stimuli. Here we examined the iPNEC and ePNEC response to potential components of the inhaled environment; the bacterial endotoxin LPS, the volatile chemical Bergamot oil, known to activate olfactory receptors on PNEC [7] and HDM extract, an allergen commonly associated with asthma. Using this approach, we identified stimuli specific changes in gene expression. Strikingly we demonstrated, for the first time, that HDM extract elicits a neuropeptide response from PNEC enriched human airway epithelium increasing release of substance P and, most markedly, inducing expression and release of CGRP.

In addition to inducing Interleukin 5 (IL-5) production in ILC2 cells, CGRP can promote a Th2 response [8] and enhance airway inflammation through Th9 cells [22]. Suppressing PNEC-derived CGRP signaling via a CGRP receptor antagonist has been shown to reduce ILC2 activation and to ameliorate asthmatic phenotypes in mice [23] suggesting a proinflammatory role for the peptide. However, another study found that sensory nerve-derived CGRP suppressed ILC2 function and allergic airway inflammation [24]. There are also studies suggesting an anti-inflammatory role for CGRP in models of allergic dermatitis [25] and airway disease [26, 27]. These contradictory findings indicate that further studies regarding the differential effects of the neuropeptide depending on factors such as source (neuronal vs non-neuronal) and timing of release in relation to allergen exposure and synergistic/antagonistic effects of co-released neuroendocrine factors, are warranted. CGRP exists in two distinct forms (αCGRP and βCGRP) encoded by separate genes (CALCA and CALCB). CALCA encodes the neuropeptide αCGRP localized in both the central and peripheral nervous system [28].

CALCB produces the neuropeptide βCGRP, present in enteric nervous system [29], and immune cells [30]. While neither CALCA or CALCB were expressed in naïve iPNEC or ePNEC cultures, primary murine and human PNECs express high levels of CALCA (95% in mouse [12] and 20% [31] to 60% [12] in human). The heterogeneity in CGRP content of human PNEC and lack of constitutive CGRP *in vitro*, might stem from the fact that *in vivo* PNEC not only produce neuropeptides, but take up and store them from the microenvironment [32]. *In vitro*, PNEC are exposed to co-cultured epithelial cells but lack input from sensory neurons present in the *in vivo* lung environment, which are a likely source of CGRP. The lack of prestored CGRP in *in vitro* cultures may explain why we did not observe CGRP release in response to Bergamot oil in contrast to a previous report by Gu et al. that demonstrated release of CGRP from *ex vivo* PNEC 15 minutes after stimulation of with Bergamot oil [7]. The lack of the ability of our model system to detect stimuli leading to rapid release of prestored CGRP is a limitation. However, we feel our observation that HDM extract can induce gene expression and subsequent CGRP release from PNEC is a significant advance in understanding the potential role of PNEC in asthma. Furthermore, the induction of CGRP production rather than immediate release is in keeping with observations that CGRP increases in both bronchoalveolar lavage and biopsies from allergic asthma patients during the late phase of allergen-induced reactions [33].

Our study further provides mechanistic insight identifying that HDM induced CGRP expression is PAR-1 mediated. The PAR1 receptor can be activated by proteases from bacteria, plants, fungi, and insects [16]. Zhang et al. previously found that HDM induced reactive oxygen species production by human airway epithelial cells was transduced through PAR1 and PAR4 [15]. CALCB induction by HDM was inhibited by both serine and cysteine protease inhibitors, suggesting multiple proteases from the HDM extract may induce CALCB expression. However we could not demonstrate CALCB induction by recombinant Der P1, a HDM component and well-described cysteine protease [17, 18] despite confirmed protease activity. Hence the nature of the HDM extract proteases that signal through PAR1 to induce CGRP warrants further investigation.

Overall, we present the first functional characterization of iPNECs and the novel ePNEC model following stimuli representative of the inhaled environment. We demonstrated, for the first time, that human PNEC respond directly to challenge HDM extract, but not LPS or volatile odorant stimuli, to produce the immunomodulatory peptide CGRP. We also provided mechanistic insight identifying that the HDM response is dependent on PAR1. This study suggests a previously unrecognized role for human PNEC in mediating a direct neuroendocrine response to allergen exposure. Taken together with recent animal model studies our results support PNEC and CGRP production by these cells as a potential therapeutic target in asthma.

### Materials and Methods iPNEC culture

Induced pluripotent stem cells (iPSCs) were purchased from Coriell Institute (□ female, 22; ◼ male, 32). The cells were maintained in mTeSR1 medium (STEMCELL Technologies) in 6-well tissue culture-treated plates on Geltrex matrix (Life Technologies). iPSC colonies were split using ReLeSR (STEMCELL Technologies). 24-well transwell inserts were coated with a combination of collagen IV (60 μg/mL) and laminin (5 μg/mL) from Sigma-Aldrich and fibronectin (5 μg/mL; Biosciences). iPSC were dissociated to a single-cell suspension using Accutase (STEMCELL Technologies). 150K iPSCs in mTeSR1 (STEMCELL Technologies) supplemented with 10 μM ROCK inhibitor Y-27632 (Selleckchem) were added to transwell inserts. Once iPSCs were 80% confluent they were differentiated through definitive endoderm, anterior foregut endoderm and lung-specific endoderm stages before they were matured to PNECs at ALI from day 17 onward using medium G (**Fig.S1 B, Table S1**) and 10 μM DAPT (Cayman Chemical) with media changes every other day [10].

### ePNEC culture

Healthy primary HBEC (● male, 52; ▲ male, 56; ○ female, 55) were purchased from ATCC (PCS-300-010). Cells were grown using the bronchial epithelial cell growth kit from ATCC (PCS-300-040). 24-well transwell inserts were coated with a combination of collagen IV (60 μg/mL) and laminin (5 μg/mL) from Sigma-Aldrichand fibronectin (5 μg/mL; Biosciences). HBEC were dissociated to a single-cell suspension using Trypsine (Sigma-Aldrich). 100K HBECs in bronchial epithelial cell growth medium were added to transwell inserts. Once HBEC were 80% confluent (1-3 days after seeding) they were matured to PNECs at ALI from day 17 onward using medium G (**Table S1, [10]**) and 10 μM DAPT (Cayman Chemical) with media changes every other day [10].

### Stimulation and inhibitor experiments

Primary HBEC, ePNEC and iPNEC were stimulated with (HDM extract (50 μl of 12, 120, 1200 and 12000 μg/mL corresponding to 0, 0.6, 6, 60, and 600 μg total protein in PBS; LoTox D. pteronyssinus antigen, Indoor biotechnologies), Lipopolysaccharide (1 ng/μl in PBS; eBioscience) and Bergamot oil (1:100 in DMSO, secondary dilution 1:1000 in PBS; Sigma-Aldrich) for 2h (RT-PCR) or 6h (ELISA). Further, PAR1 antagonist Vorapaxar was used at 80 μM and PAR1 agonist TFLLR-NH2 at 2 μM (Cayman Chemical) was added at the same time as HDM (120 μg/ml) for 2h (RT-PCR) or 6h (ELISA). Similarly, PAR2 antagonist FSLLRY (Cayman Chemical) was applied at 80 μM together with HDM (120 μg/ml) for 2h (RT-PCR) or 6h (ELISA). Recombinant Der P1 (Indoor biotechnologies) was used at 10 μM and activated with Dithiothreitol (DTT; Sigma-Aldrich) at 5mM for 20 min. Protease inhibitors used were the non-specific protease inhibitor Chymostatin, serine protease inhibitor Phenylmethylsulfonyl Fluoride (PMSF) and cysteine protease inhibitor E-64 (Sigma-Aldrich). Chymostatin (10 μg/ml), PMSF (0.25mM) and E-64 (10 μM) were pre-incubated with HDM (120 μg/ml) for 15 minutes at 37°C before that mixture was applied to ePNEC for 2h (RT-PCR) or 6h (ELISA). All experiments were carried out with technical triplicates and repeated 3 times. Detailed protocols for the following read outs of the experiments can be found in the supplementary methods: **Quantitative Real-time (qRT-PCR) analysis, Flow Cytometry, Protease activity assay, Enzyme-linked immunosorbent assay (ELISA), immunofluorescent stainings**, and **single-cell RNA sequencing**.

### Statistical analysis

Bar and dot plots were plotted in Graphpad Prism version 10.0.1 (GraphPad Software, www.graphpad.com), which was also used for statistical analysis. Normality of data distribution was assessed with the Shapiro-Wilk normality test. Statistics on normally distributed data (p>0.05)statistical were performed with one-way ANOVA and Tukey’s post -test for multiple comparisons between groups and with multiple unpaired t-tests, whereas the nonparametric Mann-Whitney (2 groups) or Kruskal Wallis (>2 groups) test were performed for not normally distributed data (Shapiro-Wilk p<0.05), *<0.05, **<0.01, ***<0.001, ****<0.0001. The test used is specified for each Figure legend. All bar graphs show mean and error bars with SEM.

## Acknowledgments

This work was funded by a grant from The Canadian Allergy Asthma and Immunology Foundation (CAAIF) and the Canadian Institute for Health Research (CIHR-IRSC:0756000040). PF is the AstraZeneca (Canada) Chair in Asthma and Obstructive Lung Disease.

