## Supplementary figures and images for "Human pulmonary neuroendocrine cells respond to House dust mite extract with PAR-1 dependent release of CGRP"

### Fig. S1

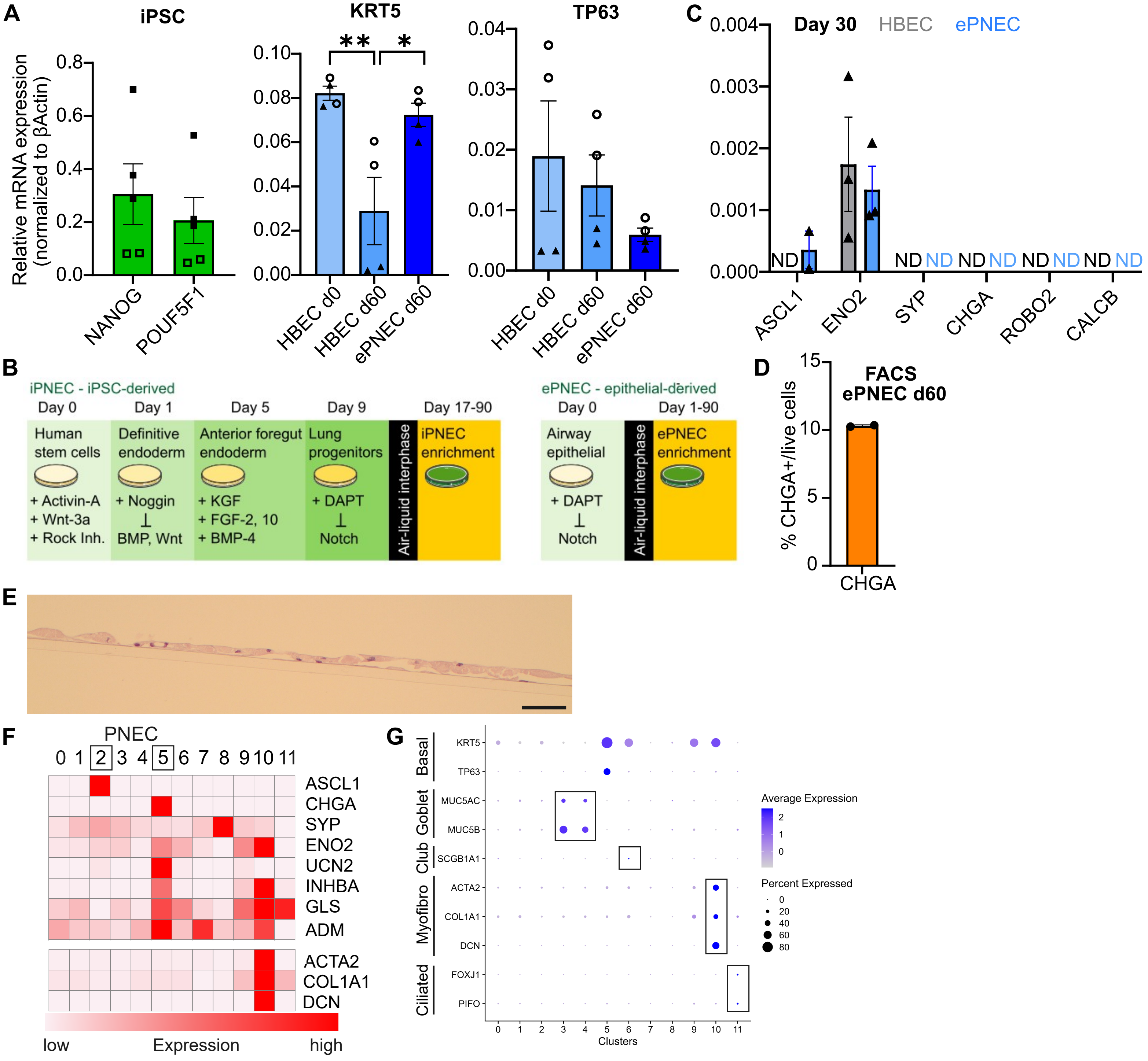

### Fig. S2

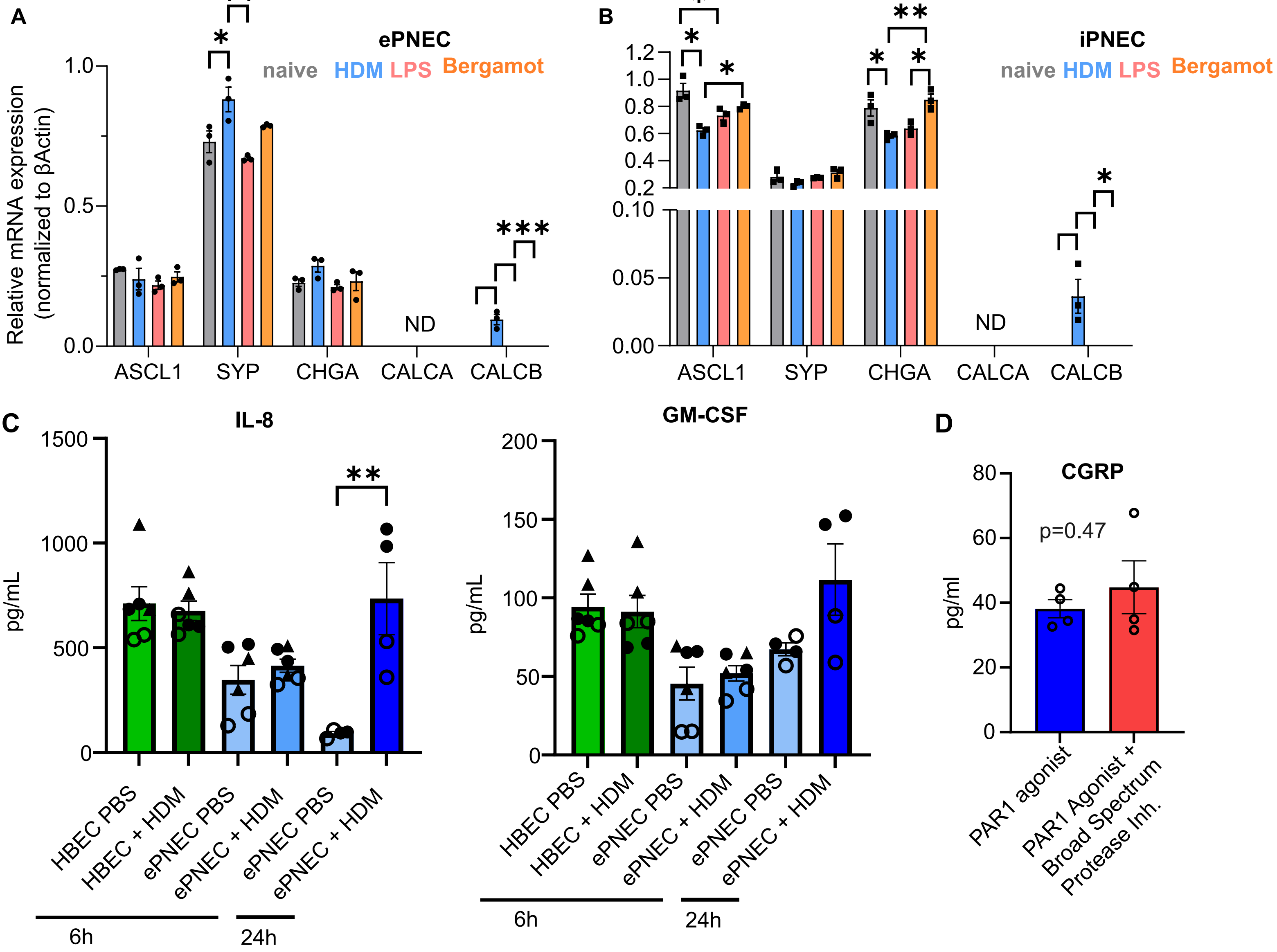

### Fig. S3

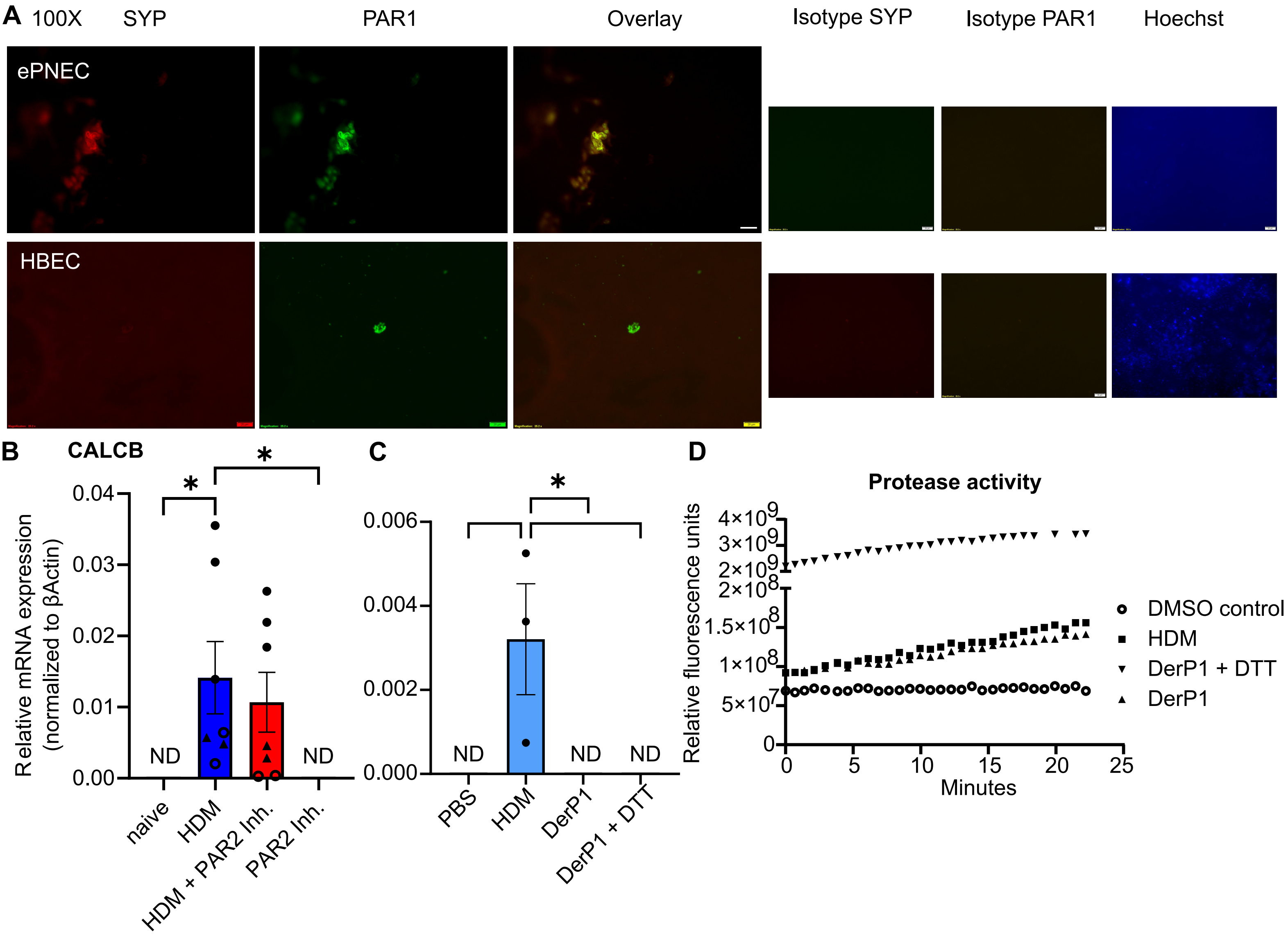
