## Supplement for "Human pulmonary neuroendocrine cells respond to House dust mite extract with PAR-1 dependent release of CGRP"

### Supplementary Materials and Methods

#### Real-time (RT-PCR) analysis

Total RNA was extracted using TRIzol reagent according to the manufacturer's instructions (Thermo Fisher). The purity and concentration of total RNA were determined by Nanodrop spectrophotometer (ND-1000, Thermo Fisher), and total RNA was reverse transcribed using the SuperScript III Platinum Two-Step qRT-PCR kit (Thermo Fisher Scientific). Amplifications for 10 µl quantitative polymerase chain reaction mixture (5 µl SYBR green, 10 ng DNA and forward and reverse primers at 300nM) were performed on the CFX96 Real-Time C1000 Thermo Cycler (Bio-Rad) using the following parameters: Initial denaturation at 95°C for 2 minutes, denaturation at 95°C for 15 seconds, annealing at 60°C for 1 minute, and extension at 72°C for 1 minute (40 cycles). The $2^{-\Delta\Delta CT}$ method was used to analyze the relative changes in gene expression. The primers were purchased from Thermo Fisher and the sequences designed can be found in **Table S2**.

#### Flow Cytometry

Day 60 differentiated ePNEC cultures were dissociated for 2 min. in trypsin at 37°C to obtain single-cell suspensions. Cells were then washed in 200µL PBS and stained with the live/dead cell marker ef780 (1:2500 in PBS, Thermo Fisher) for 15 min. at room temperature. Next, cells were washed in FACS buffer (PBS, 2 mM EDTA + 2% (v/v) FBS), blocked with Fc block (1:50 in FACS buffer) for 10 min. on ice, fixed in 100µL BD Cytofix for 20 min. on ice, washed in 100µL Perm/wash two times and stained with anti-human CHGA-AF488 conjugated antibody (1:50 in Perm/wash; clone NB120, NovusBio) for 25 min. on ice. Finally, cells were washed in Perm/wash and resuspended in 200µLFACS buffer. Flow cytometry data were collected on a BD LSR Fortessa-II (BD Bioscience, California) using FACS Diva software (BD Bioscience, California). Results were analyzed using FlowJo (TreeStar, Ashland, Oregon).

#### Protease activity assay

The fluorogenic substrate Boc-Gln-Ala-Arg-AMC (Enzo Biochem) was used for measuring protease activity in HDM at 120 µg/ml and Der P1 at 10 µM with and without DTT activation at 5mM for 20 min. at RT. The assay was conducted in a total volume of 200 µL which contained 100 µL of samples, 5 µL of a 5mM stock of the substrate and 95 µL of 100mM Tris HCl (pH 8.5) in 96-well plates. DMSO and Trypsin served as negative and positive controls, respectively. Relative fluorescence was determined with excitation at 380 nm and emission at 460 nm over a period of 30 min. and measurements every 46 sec. Protease activity was determined as the slope of the fitted curve plotting relative fluorescent units against time.

**ELISAs**

Frozen supernatant samples were thawed and subjected to analysis using the following Enzyme-linked immunosorbent assay (ELISA) kits: Enolase (DENL20, R&D Systems Inc.), CGRP ELISA kit (LS-F37468, LS Bio) and Substance P (KGE007, R&D Systems Inc.) following each manufacturer's protocol. Standard curves were used for linear regression to determine protein concentrations. Media and HDM (1200 µg/ml) was used as a negative control. All experiments were carried out with experimental triplicates. GM-CSF and IL-8 were measured by Eve Technologies Corp. (Calgary, Canada) according to a manufacturer's protocol by MilliporeSigma (Burlington, USA).

**Immunofluorescent stainings**

Transwells containing cells were washed 3X with room temperature PBS for 5 minutes each and fixed in methanol overnight at -20°C. The next morning, methanol was removed and the bottom portion of the inserts was dipped int -20°C acetone for 1 minute. Each insert was washed 3X with PBS for 5 minutes each and incubated with CAS block (Thermo Fisher Scientific) at room temperature for 1 hour. Primary antibody was diluted in CAS block and 200 µL of the diluted mixture was added to the apical compartment and 500 µL to the basal compartment. Plates were incubated overnight at 4C and washed 3X with PBS for 5 minutes each the next morning. Relevant secondary antibodies were diluted at 1:500 in CAS buffer and applied together with Hoechst stain for nuclei (ThermoFisher Scientific) diluted at 1:5000 to the transwells at room temperature for 2 hours in the dark (full list of antibodies in **Table S3**). Next, inserts were washed 3X with PBS for 5 minutes each and the polyester membranes were cut out using surgical scissors to be mounted using SlowFade Diamond Antifade mountant (Thermo Fisher Scientific). The specificity of immunostaining was verified via isotype controls. All images were collected on an Olympus IX81 epifluorescence microscope and analyzed using cellSens software (Olympus Canada). All experiments were carried out with technical triplicates and repeated 3 times.

**Single cell RNA sequencing**

Day 60 differentiated ePNEC cultures were dissociated for 2 min. in trypsin at 37°C to obtain single-cell suspensions. Cells were frozen in 50% FBS, 10% DMSO and 40% medium. The following was performed with Active Motif: Libraries were performed according to 10x manufacturing instructions. Briefly, we utilized a Countess to measure the amount of cells within the sample. Cells were resuspended in a master mix and loaded together with partitioning oil and gel beads. This mixture was loaded onto Chip G to generate gel bead-in-emulsion. The poly-A RNA from the cells loaded into the chromium controller was then retrotranscribed into cDNA which contained a UMI, a 10x barcode, and an Illumina R1 primer sequence. The cDNA was cleaned up with Silane DynaBeads. We then performed a quality control check for the quality of the full-length cDNA by tapestation. Afterwards, full-length cDNA was fragmentated and ligation occurred to add the Illumina TruSeq adaptor. PCR occur afterwards to add i5 and i7 indexes as well as P5 and P7 sequences so that the sample could be sequence on an Illumina sequencer. Final cDNA libraries were measured by Tapestation and then sequenced on a NovaSeq for 30,000 read pairs per cell. Data is available on Genbank (SRR29297737, BioProject PRJNA1120168).

**Single cell RNA sequencing data analysis**

High quality cellular barcodes were selected using a mixture of both adaptive and selective thresholds. To pass quality control, a cellular barcode must match all three of the following criteria: 1) The number of unique molecular identifiers (UMIs) must be within three median absolute deviations (MADs) of the population median. 2) The number of expressed genes must be within three MADs of the population median. 3) The percentage of reads mapping to mitochondrial genes must be under 15%. To determine the optimal cluster resolution, the clustering-tree method was applied. The cluster tree provides an overview of the relationship between clusters at multiple resolutions and facilitates the selection of an optimal resolution by incorporating meta-information. The cluster tree plot consists of multiple layers of clustered nodes, where each layer represents a different resolution, and edges represent the transition of clusters between resolutions. The edges are color-coded based on the number of cells they encompass, with transparency adjusted using the in-proportion metric to emphasize more significant edges. By default, the size of nodes is adjusted according to the number of cells in the cluster, and their color indicates the clustering resolution.

The stability index calculated using the SC3 package [34] measures the stability of clusters across different resolutions. To determine the ideal cluster resolution, we first calculated the average stability index value for each resolution and chose the resolution that yields the highest stability value. Seurat (V4) was applied to downstream analysis. After a quality control step, we employed a global-scaling normalization method "LogNormalize" that normalizes the feature expression measurements for each cell by the total expression, multiplies this by a scale factor 10000 via function NormalizeData in Seurat. Then top 2000 highly variable genes (HVGs) were selected for initial clustering of cells. Scaled data was calculated through Seurat ScaleData function which applies Linear transformation (a standard pre-processing step prior to dimensional reduction techniques). To perform dimension reduction, we first applied RunPCA with 30 PCs. UMAP embedding of cells were performed via RunUMAP with 30 dimensions. For clustering, we used the function FindClusters that implements Shared Nearest Neighbor modularity optimization-based clustering algorithm on 20 PC components with the optimal resolution predicted in the previous step. Differential analysis was performed using the FindAllMarkers function with the following parameters: 1) Wilcoxon model, 2) this test is only performed for genes that are expressed in at least 10% of the cells in in either of the two populations. PNEC clusters were identified by exclusive expression of CHGA or ASCL1. Markers for identification of the other cell types can be found in **Table S4** and **Fig.S1 F, G**.

#### Re-analysis of publicly available data

PNEC expression of receptors implicated in allergen sensing was conducted by a re-analysis of published single cell transcriptomics data of iPSC derived PNECs (GSE146990, [16]) using Cellenics hosted by Biomage (https://biomage.net/).

### Supplementary Figures

**Fig. S1 Characterization of undifferentiated iPSCs and HBECs and differentiated iPNECs and ePNECs.** **A** iPSC (□ female, 22; ■ male, 32) and primary HBEC (▲ male, 56; ○ female, 55) characteristic marker expression shown with qRT-PCR. **B** Graphic illustration of iPSC to iPNEC and HBEC to ePNEC differentiation. **C** qRT-PCR of selected PNEC-related genes for HBEC and ePNEC at day 30 of culture (male, 56). **D** Co-staining of SYP and CHGA in iPNECs. RT-PCR data are normalized to internal control β-actin. **E** Heatmap showing expression of typical neuroendocrine markers for identification of ePNEC clusters from sc-RNA-Seq. Color scale shows relative expression within a row to highlight cluster differences. **G** Dot plot showing genes used for identification of co-cultured cells from scRNA-Seq. Data shown for mean ± SEM. One-way ANOVA for **A**, Mann-Whitney test for **C**. *<0.05, **<0.01, ***<0.001, ****<0.0001. Abbreviations: BMP4 – Bone morphogenetic protein 4; CHGA – Chromogranin A ; DAPT - 3tert-Butyl(2S)-2-((2-(3,5-diflurophenyl) acetyl) amino) propanoyl)amino)-2-phenylacetate; ePNEC – epithelial-derived pulmonary neuroendocrine cells; FGF – Fibroblast growth factor;  Inh. – Inhibitor; iPNEC – iPSC-derived pulmonary neuroendocrine cells; iPSC – induced pluripotent stem cells; KGF – Keratinocyte growth factor; KRT5 – Keratin 5; NANOG – Nanog Homeobox; POU5F1 – POU Class 5 Homeobox1; SYP – Synaptophysin; TP63 – Tumor protein p63.

**Fig. S2 Canonical marker expression and CGRP ELISA after HDM stimulation.** **A, B** qRT-PCR of canonical PNEC markers ASCL1, SYP, CHGA, CALCA and CALCB after 2h stimulation with LPS (1ng/µl), Bergamot oil (1:1000, PBS) and HDM (60 µg total protein in PBS) in ePNEC (male, 52) (**A**) and iPNEC (male, 32) (**B**). **C** GM-CSF and IL-8 protein levels of ePNEC and HBEC for 6h (● male, 52; ▲ male, 56; ○ female, 55) and 24h (● male, 52; ○ female, 55) HDM stimulation (1200 µg/ml in PBS), respectively. **D** CGRP release after 6h PAR1 agonist (TFLLR-$NH_2$ at 2 µM in PBS) stimulation from ePNEC (○ female, 55) with and without addition of a broad spectrum protease inhibitor (Chymostatin 10 µg/ml) to the basolateral chamber simultaneous to PAR1 agonist exposure. Each dot represents one well and data is shown in mean ± SEM. One-way ANOVA for **A, B**; Kruskal-Wallis test for **C;** unpaired t-test for **C**, **D**. *<0.05, **<0.01, ***<0.001, ****<0.0001. Abbreviations: ASCL1 – Achaete-Scute Family BHLH Transcription Factor 1; CALCA – Calcitonin-related polypeptide alpha; CALCB – Calcitonin-related polypeptide beta;; CHGA – Chromogranin A; ePNEC – epithelial cell derived pulmonary neuroendocrine cells; HDM – House dust mite; iPNEC – iPSC-derived pulmonary neuroendocrine cells;  iPSC – induced pluripotent stem cells; LPS – Lipopolysaccharide; SYP – Synaptophysin.

**Fig. S3 Mechanistic insight into HDM-induced CGRP expression.** **A** SYP and PAR1 immunohistochemical co-staining in naive ePNEC and control HBEC (male, 56) and isotype controls. 40X magnification and scale bars at 20 µm. **B** qRT-PCR of CALCB in control (PBS), 2h HDM (1200 µg/ml equaling 60 µg total protein in PBS), HDM and PAR2 antagonist FSLLRY (80 µM in PBS) co-incubation or FSLLRY presence alone  in ePNEC (● male, 52; ▲ male, 56; ○ female, 55) at d60 of culture. **C** qRT-PCR of CALCB in ePNEC for control (PBS), 2h HDM (1200 µg/ml), recombinant Der P1 with and without DTT activation. **D** Protease activity assay using Boc-Gln-Ala-Arg-AMC for assessing HDM and Der P1 with and without DTT activation. Each dot represents one well and data shown for mean ± SEM. One-way ANOVA for **B**. *<0.05, **<0.01, ***<0.001, ****<0.0001. Abbreviations: ASCL1 - Achaete-Scute Family BHLH Transcription Factor 1; Avg. – Average; CHGA – Chromogranin A ; DTT – Dithiothreitol; ENO2 – Enolase 2; Exp. – Expression; F2R – Coagulation Factor II Thrombin receptor; F2RL1 - Coagulation Factor II Thrombin receptor like 1; F2RL2 - Coagulation Factor II Thrombin receptor like2; F2RL3 - Coagulation Factor II Thrombin receptor like 3; PNEC – Pulmonary neuroendocrine cells; SYP – Synaptophysin.

### Supplementary Tables

**Table S1: Media used for differentiation of iPNEC and ePNEC.**

| ***Reagent*** | ***Final concentration*** | ***Supplier*** |
|---|---|---|
| **Medium B** | | |
| RPMI 1640 | | Thermo Fisher Scientific, Waltham, MA, USA |
| Wnt-3a | 25 ng/mL | R&D Systems Inc., MN, USA |
| Activin A | 100 ng/mL | Proteintech Group, IL, USA |
| **Medium C** | | |
| RPMI 1640 | | |
| FBS | 1% | Corning, NY, USA |
| Activin A | 100 ng/mL | Proteintech Group, IL, USA |
| **cSFDM medium** | | |
| IMDM | 75% | Thermo Fisher Scientific, Waltham, MA, USA |
| Ham’s F12 | 25% | |
| Glutamax | 1 X | |
| B-27 with RA | 0.5 X | |
| N-2 | 0.5 X | |
| BSA | 0.75% (w/v) | Sigma-Aldrich, MO, USA |
| Ascorbic Acid | 50 µg/mL | |
| 1-Thioglycerol | 4.5 x $10^{-4}$ M | |
| **Medium D** | | |
| cSFDM | | |
| SB431542 | 500 nM | Selleck Chemicals, TX, USA |
| Noggin | 20 ng/mL | Pepro Tech, NJ, USA |
| **Medium E** | | |
| cSFDM | | |
| SB431542 | 500 nM | Selleck Chemicals, TX, USA |
| BMP-4 | 5 ng/mL | Proteintech Group, IL, USA |
| FBS | 2% | Corning, NY, USA |
| **Medium F** | | |
| cSFDM | | |
| BMP-4 | 5 ng/mL | Proteintech Group, IL, USA |

| | | |
|---|---|---|
| FGF-2 | 10 ng/mL | Thermo Fisher Scientific, Waltham, MA, USA |
| FGF-10 | 10 ng/mL | Pepro Tech, NJ, USA |
| KGF | 10 ng/mL | |
| **Medium G** | | |
| DMEM/F-12 | | Thermo Fisher Scientific, Waltham, MA, USA |
| Ultroser G | 2% | Crescent Chemical, NY, USA |
| Fetal Clone II | 2% | Hyclone, UT, USA |
| Insulin | 2.5 µg/mL | Lonza, OR, USA |
| Bovine Brain Extract | 22.5 µg/mL | |
| Transferrin | 2.5 µg/mL | Sigma-Aldrich, MO, USA |
| Hydrocortisone | 20 nM | |
| 3,3‘,5-Triiodo-L-thyronine | 500 nM | |
| Epinephrine | 1.5 µM | |
| Retinoic Acid | 10 nM | |
| Phosphoryl ethanolamine | 250 nM | |
| Ethnaolamine | 250 nM | |
| DAPT | 10 µM | Cayman Chemical, Ann Arbor, MI, USA |

**Table S2**: Quantitative RT-PCR primers.

| Gene | Forward | Reverse |
|---|---|---|
| ADCYAP1 | 5′-CCACTCGGACGGGATCTTC-3′ | 5′-GCCGCCAAGTATTTCTTGACAG-3′ |
| ASCL1 | 5′-CCCAAGCAAGTCAAGCGACA-3′ | 5′-AAGCCGCTGAAGTTGAGCC-3′ |
| CALCA | 5′-ATGGGCTTCCAAAAGTTCTCC-3′ | 5′-GCCGATGAGTCACACAGGTG-3′ |
| CALCB | 5′-ACCCGGCCACACTCAGTAA-3′ | 5′-GGGCACGAAGTTGCTCTTCA-3′ |
| CCDC25 | 5′-CACCTTGTGAAGGCCAATAGC-3′ | 5′-CTGCCCCACATCCATGTCAG-3′ |
| CHAT | 5′-CAGCCCTGCCGTGATCTTT-3′ | 5′-TGTAGCTGAGTACACCAGAGATG-3′ |
| CHGA | 5′-TAAAGGGGATACCGAGGTGATG-3′ | 5′-TCGGAGTGTCTCAAAACATTCC-3′ |
| CRH | 5′-GGGAACCTCAACAAGAGCCC-3′ | 5′-AACACGCGGAAAAAGTTGGC-3′ |
| CXCL13 | 5′-GCTTGAGGTGTAGATGTGTCC-3′ | 5′-CCCACGGGGCAAGATTTGAA-3′ |
| DDC | 5′-TGGGGACCACAACATGCTG-3′ | 5′-TCAGGGCAGATGAATGCACTG-3′ |
| ENO2 | 5′-CTGATGCTGGAGTTGGATGG-3′ | 5′-CCATTGATCACGTTGAAGGC-3′ |
| GAD67 | 5′-GCTTCCGGCTAAGAACGGT-3′ | 5′-TTGCGGACATAGTTGAGGAGT-3′ |
| GRP | 5′-AAAGAGCACAGGGGAGTCTTC-3′ | 5′-TCCTTTGCTTCTATGAGACCCA-3′ |
| KRT5 | 5′-CCAAGGTTGATGCACTGATGG-3′ | 5′-TGTCAGAGACATGCGTCTGC-3′ |
| NANOG | 5′-TTTGTGGGCCTGAAGAAAACT-3′ | 5′-AGGGCTGTCCTGAATAAGCAG-3′ |
| OR2W1 | 5′-ACACTCACTCTGAATTTGCCC-3′ | 5′-TTTCAACTGTTGTGGTGTCTACA-3′ |
| PAR1 | 5′-GTATCCCATGCAGTCCCTCTCC-3′ | 5′-GTAATGCGCAATCAGGAGGACG-3′ |
| PENK | 5′-TCTGAACCCGGCTTTTCCAA-3′ | 5′-TACGCAAGCCAGGAAGTTGAT-3′ |
| POUF5F1 | 5′-CTGGGTTGATCCTCGGACCT-3′ | 5′-CCATCGGAGTTGCTCTCCA-3′ |
| PPT1 | 5′-TGTTTTTGGACTCCCTCGATG-3′ | 5′-CATGCCAGTATTCGGCTTGC-3′ |

| | | |
|---|---|---|
| ROBO2 | 5′-GTTTGTGTTGCGAGGAACTATCT-3′ | 5′-GTTTTGTCGGAAGTCATCTCGTA-3′ |
| SST | 5′-CCCAGACTCCGTCAGTTTCT-3′ | 5′-AAGTACTTGGCCAGTTCCTGC-3′ |
| SV2 | 5′-TAGACCAGGCACTCATTGGC-3′ | 5′-ACCCCTCCCCACAGTTACTT-3′ |
| SYP | 5′-CTCAGCATCGAGGTCGAGTTC-3′ | 5′-GAGGAGTAGTCCCCAACTAAGAA-3′ |
| SYT1 | 5′-GCTGCTGGTAGGGATCATTCA-3′ | 5′-GTTTTTCGGTGGACTTTTGTCTC-3′ |
| TP63 | 5′-GGACCAGCAGATTCAGAACGG-3 | 5′-AGGACACGTCGAAACTGTGC-3′ |
| TUBB3 | 5′-GGCCAAGGGTCACTACACG-3′ | 5′-GCAGTCGCAGTTTTCACACTC-3′ |
| β-Actin | 5′-CATGTACGTTGGTATCGAGGC-3′ | 5′-CTCCTTAATGTCACGCACGAT-3′ |

**Table S3**: Primary and secondary antibodies used.

| **Name** | **Brand** | **Catalog number** | **Dilutions** |
|---|---|---|---|
| SYP | Abcam | ab32127 | 1:300 |
| ENO2 | Santa Cruz Biotechnology | Sc-271384 (A-5) | 1:100 |
| CHGA | Proteintech | 60135-1-Ig | |
| PAR1 | Novus Biologicals | NBP1-71770SS (N2-11) | 1:100 |
| IgG (H+L) AF488 | Fisher Scientific | A21202 | 1:500 |
| IgG (H+L) AF555 | Fisher Scientific | A31570 | 1:500 |
| IgG (H+L) AF555 | Fisher Scientific | A31572 | 1:500 |

**Table S4** Markers used for identification of clusters from single cell RNA sequencing.

| **Cell type** | **Markers** |
|---|---|
| PNEC | CHGA or ASCL1 (exclusively expressed in identified clusters), ENO2, UCN2, INHBA, GLS, ADM |
| Goblet cells | MUC5AC, MUC5B |
| Club cells | SCGB1A1 |
| Basal cells | NGFR, KRT5, TP63 |
| Ciliated epithelial cells | FOXJ1, PIFO |
| Myofibroblasts | ACTA2, VIM, COL1A1, DCN |
